# Refining Protein Co-Membership in Reactome: A Weighted Scoring Approach Based on Hierarchy and Sequences of Patwhays

**DOI:** 10.64898/2026.09.30.755656

**Authors:** Cécile Beust, Olivier Dameron, Nathalie Théret, Emmanuelle Becker

## Abstract

In network biology, analyzing pathways is critical to study mechanisms or sequences of biological events. The pathway co-membership approach aims to link proteins belonging to the same pathway inside a network that is a clique. Although this method is great for projecting pathway knowledge onto proteins, it remains unsuitable for some downstream applications relying on graph distances like random walks with restart. To address this limitation, we introduce a weighted pathway co-membership method that incorporates biological distances between proteins. We achieve this by developing an abstraction framework for Reactome pathways, which captures both their hierarchical organization and sequential relationships. By assigning weights to these abstractions, we define a scoring system for proteins within the pathway co-membership clique. We illustrate our approach using a Reactome pathway (“Signaling by TGF-beta receptor complex”, `R-HSA-170834`), and explain how we generate the weighted co-membership cliques of the 29 Reactome top-level pathways. Next, we construct the entire weighted pathway co-membership network of the Reactome knowledge base. Validation and evaluation of our method is performed using cancer hallmark gene sets and a random walk with restart experiment, comparing our approach to state-of-the-art methods. We show that our approach outperforms state-of-the-art methods for gene prioritization on cancer hallmarks.

**Code and Data availability:** SPARQL queries are available on GitHub and are integrated within a Jupyter notebook^*^. The weighted co-membership cliques of the 29 Reactome top-level pathways are available on NDEx^†^.

**Graphical Abstract:** 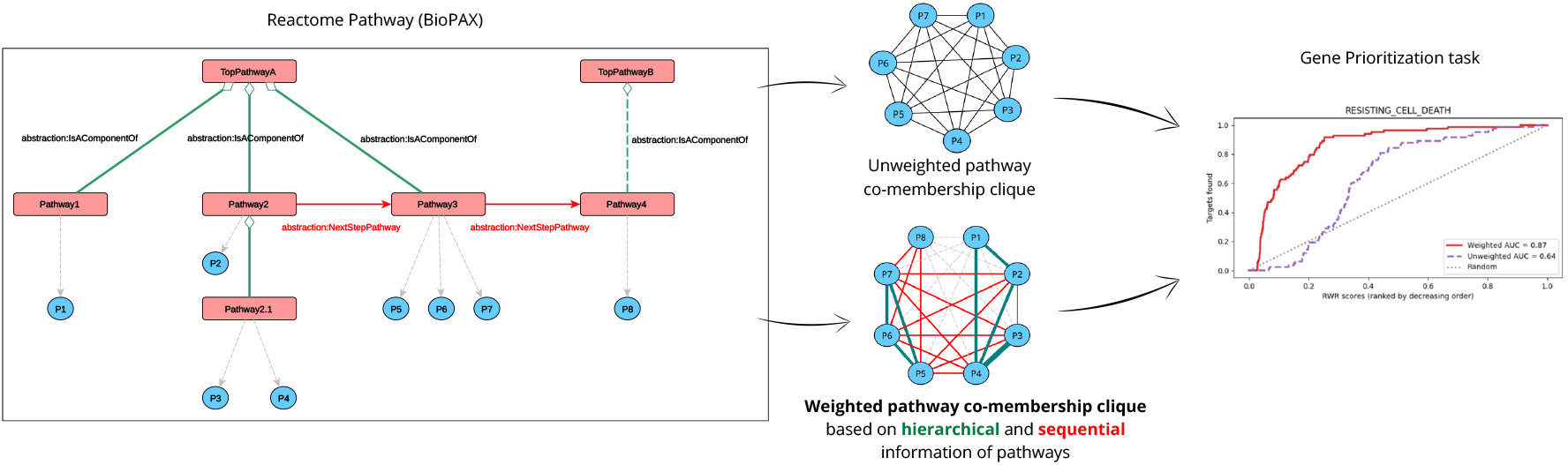

**Key Messages:**

- We revisit methods to score protein interactions based on their pathway co-membership by leveraging knowledge on the composition and sequences of Reactome pathways
- We show that our approach outperforms state-of-the-art methods for gene prioritization on cancer hallmarks
- We pre-computed and deposited on NDEx the weighted pathway co-membership cliques of the 29 top-level pathways of Reactome independently^‡^, and on the entire Reactome knowledge base to facilitate exploration of protein interactions.

## Introduction

Interactions between proteins are fundamental in biological processes, and their understanding is key for gaining insights into biological mechanisms. To decipher these mechanisms, it is essential to characterize protein interactions not merely as isolated events, but as components of larger, organized functional units. Biological pathways, which are series of molecular interactions leading to a certain product or change in the cell^1^, serve as critical functional modules for characterizing protein interactions (Nayar and Altman, 2024). By analyzing pathways as structured networks of interacting proteins, researchers can move beyond simple gene lists to capture the systematic logic of cellular functions. Methods such as the graphite R package (Sales et al., 2012) propose ways to abstract pathway topology into molecular interaction networks, by taking into account the topological complexity of the pathways (protein complexes, gene families, signal propagation through chemical compounds, etc). Such development emphasizes the crucial need of leveraging pathway topology information for accurate interpretation of protein interaction networks.

In this paper, **we introduce a weighted version of the pathway co-membership approach that takes into account the biological distance of proteins in the pathway co-membership clique**. We tested our approach on Reactome pathways (Fabregat et al., 2018), since Reactome is a comprehensive resource of biological pathways (Bora et al., 2026; Beust et al., 2024). In addition, Reactome can be exported in the BioPAX exchange format that standardizes pathway information into a knowledge graph. We defined a scoring system of Reactome pathways and their interactions following an **abstraction strategy that extracts both the hierarchy of the pathways and their sequentiality** from the BioPAX graph. The scoring of Reactome pathways and their relations allow to score pairs of proteins by getting back their pathway parents and computing distances between them. We evaluated the benefits of taking into account compositional and sequential relations versus classical co-membership methods using gene prioritization tasks on various cancer hallmarks gene sets, and showed that our method outperforms state-of-the-art methods for this specific task.

### State of the art

It is important to highlight the difference between weighting relations between pathways and weighting relations between proteins depending on their pathways: in the first case we can build the network where the nodes are the pathways and the interactions correspond to relations between pathways, and in the second case the nodes are the proteins/genes and the interactions correspond to pathway co-membership.

Weighting relations between pathways was introduced to explore visually pathway databases or to capture relations between pathway’s annotations of different knowledge bases. In this case it addresses the question *“to what extent the pathways involve the same participants (e*.*g. proteins, genes, molecules*…*)?”*. This can be measured using metrics such as Jaccard index (Parikh et al., 2012) or overlap coefficients (Domingo-Fernández et al., 2019). These metrics are normalized and can be used to work simultaneously with different databases such as KEGG (Kanehisa and Goto, 2000), Reactome (Fabregat et al., 2018), WikiPathways (Pico et al., 2008; Agrawal et al., 2024) and GO BP (Ashburner et al., 2000). These metrics allow to indirectly take into account the information about the composition hierarchy of pathways if the participants of a pathway also include the participants of its sub-pathways. Pathway specificity is taken into account with a normalization as a function of pathways size. Note that these metrics focus on the compositional similarity of pathways, which is different from functional similarity. For example, two pathways can have disjoint sets of participants while being functionally related as sub-pathways of the same super-pathway.

Weighting relations between proteins as a function of the pathways they participate in was introduced to answer a different question: *“to what extent proteins are involved in common pathways ?”*. Proteins can be represented as vectors (often binary) of the pathways they participate in, thus allowing to take into account pathway compositional hierarchy. These vectors are then compared according to different families of functions. A simple approach consists in counting the number of common pathways between two proteins, and eventually applying a threshold. Other approaches rely on different strategies to normalize this number of common pathways, either using a Jaccard index (Jothi et al., 2007) or Cosine similarity (Wu et al., 2015). Note that the Jaccard index can be used for both comparing pathways according to their proteins (as seen previously) as well as comparing proteins according to the pathways they participate in. While in the first case the Jaccard index depend on the sizes of the pathways, this is not true for the second case where it depends on the total number of pathways and the hierarchy structure.

If these approaches properly answer the question of belonging to commons pathways, they are not meant to answer the question “to what extent are the proteins functionally related ?” because (i) they are sensitive to unbalanced pathway compositional hierarchies and (ii) they do not take into account pathway sizes thus scoring equally general pathways and specific ones. Indeed, functional similarity between pathway participants is inversely proportional to the number of participants of the pathway. As an illustration, two proteins among the 2, 216 participating in the “Metabolism” pathway (`R-HSA-1430728`) are less functionally related than two proteins among the 38 involved in the “Triglycerides catabolism” pathway (`R-HSA-163560`). To overcome these limitations, strategies can take into account information content and semantic similarity measures (for a review see (Pesquita, 2017)). Most approaches leverage the biological process hierarchy of the Gene Ontology, and several semantic similarity measures have been proposed allowing to take into account the specificity of pathways through their compositional hierarchy.

## Materials and Methods

### Materials

We choose to work on the BioPAX exports of Reactome pathways, since this language allows to finely describe pathways using the rich vocabulary of the BioPAX ontology.

We used the version 96 of the Reactome knowledge base (April 2026), and downloaded the BioPAX export^2^. We selected the *Homo sapiens* BioPAX files within the downloaded folder.

For experiments involving individual Reactome pathways or top-level pathways (i.e. super-pathways corresponding to main biological processes), individual BioPAX files were downloaded from Reactome.

For executing SPARQL queries, we loaded BioPAX files into an Apache Jena Fuseki endpoint^3^.

### Requirements for a scoring system of pathway co-membership interactions among proteins

For weighting pathway co-membership interactions among proteins, our metric respects some requirements:

- Principle 1: There is a co-membership relation between each pair of proteins of the same pathway. For a given pathway, it creates a maximal clique between its participants.
- Principle 2: Co-membership is a weighted relation whose weight is a monotonous decreasing function of the size of the most informative common pathway: the smaller the common pathway, the greater the weight.
- Principle 3: Proteins can participate in multiple pathways. Thus, two proteins p1 and p2 can be members of several common pathways. The overall co-membership score of p1 and p2 is a function of all the scores according to these common pathways.
- Principle 4: `bp3:nextStep` relations between pathway steps are taken into account. Two proteins participating in such sequence of steps are indeed functionally-linked and are considered as members of a common (possibly unspecified) pathway.

These principles are illustrated in Supplementary Figure S-1.

### Abstracting the hierarchy of Reactome pathways

Starting from the BioPAX model of a Reactome pathway, we extracted its compositional hierarchy using a SPARQL query (Figure 1) using the `bp3:pathwayComponent` relationships. In our abstracted graph, the compositional hierarchy of pathways is modeled by the `abstraction:IsAComponentOf` relationships.

**Figure 1.**
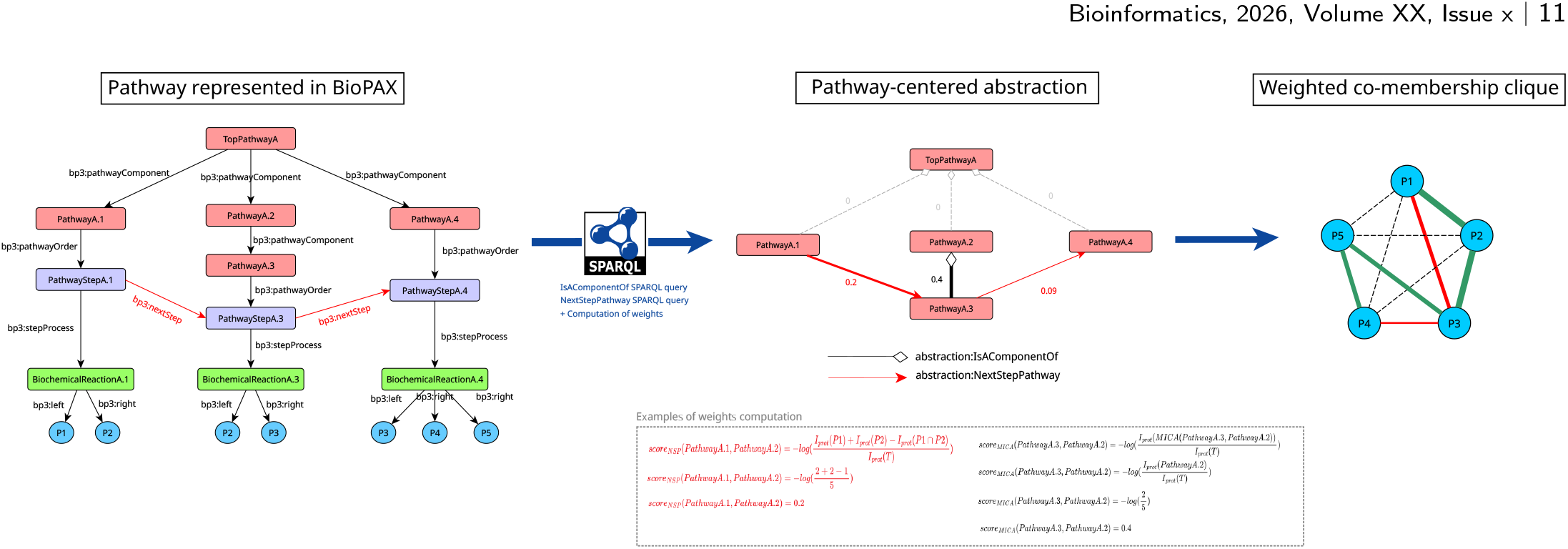
Abstracting BioPAX pathways. A toy example of a pathway represented in BioPAX is shown on the left part of the figure. In this BioPAX graph, instances of `bp3:Pathway` are represented as red nodes. Their `bp3:PathwaySteps` are represented in purple. The associated instances of `bp3:BiochemicalReaction` and `bp3:Protein are` represented in green and blue respectively. The BioPAX relationships `bp3:pathwayComponent` are those describing the hierarchy of pathways. Also, `bp3:nextStep sequence` links `PathwayStepA.1 to PathwayStepA.3` and `PathwayStepA.3 to PathwayStepA.4`. This BioPAX graph is abstracted into a simple graph presented on the right part of the figure, thanks to SPARQL queries. The abstracted graph presents two types of weighted edge types, `abstraction:IsAComponentOf edges` (dotted lines) for pathway composition relationships and `abstraction:NextStepPathway` edges (red lines) for pathway sequences.

We introduced edge weights in our pathway abstraction method, in order to reflect as much as possible the biological distance between pathways in the graph. We used the notion of information content (IC) to define our metric. The information content in an ontology measures how specific is a concept (Pesquita, 2017; Seco et al., 2004). First, pathway nodes are scored with regard to their information content in proteins (*IC*_*prot*_) derived from the Resnik information content formula (Resnik, 1995):

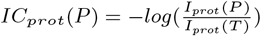

Where *P* is a pathway in the abstraction graph, *I*_*prot*_(*P* ) is the number of proteins annotated by pathway *P*, and *I*_*prot*_(*T* ) is the total number of proteins annotated by all pathways in the graph. The proteins counted are identified by UniProt identifiers (Pundir et al., 2017) found in the BioPAX model. `abstraction:IsAComponentOf` edges are scored using the information content in proteins of the most informative common ancestor (MICA) of the two pathways. This score is called *score*_*MICA*_ and is computed as defined below:

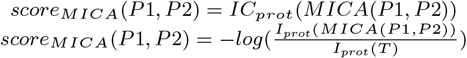

Here, *P*1 and *P*2 are two pathways connected by a `abstraction:IsAComponentOf` edge. *MICA*(*P*1, *P*2) is the pathway that is the most informative common ancestor of *P*1 and *P*2.

### Abstracting the sequences of Reactome pathways

We extracted the sequentiality of the pathways in the BioPAX graph following the BioPAX relationship `bp3:nextStep` between pathway steps. In BioPAX, instances of `bp3:Pathway` are composed of instances of `bp3:PathwayStep` that correspond to either sub-pathways or associated biochemical reactions and catalysis events. These pathway steps can be connected using `bp3:nextStep` relationships. We abstracted this by extracting the `bp3:nextStep` sequences between pathway steps and reporting it to pathways (Figure 1).

`abstraction:NextStepPathway` (NSP) edges are scored using a similar information content in protein system. For this type of edges, we do not use the MICA of pathways, but consider the direct union of their annotated proteins:

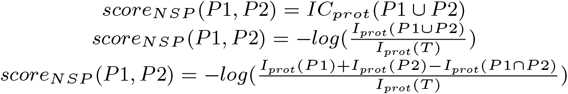

Where *P*1 and *P*2 are two pathways of the abstraction linked by a `abstraction:NextStepPathway` edge.

### Defining a scoring system for protein co-membership interactions in a pathway

We use the precomputed abstraction of the pathway in order to generate the weighted clique of protein pathway co-membership interactions as illustrated in Figure 3. First, we get back the set of proteins annotated by each pathway in the abstraction and compute two matrices of scores between pairs of proteins:

- The matrix of **MICA scores** gathers, for all protein pairs in the graph, the value of the information content in proteins of the MICA of their two pathway parents.
- The matrix of **NSP scores** (NextStepPathway scores) gathers, for all protein pairs in the graph, the highest NSP score between their respective pathways, including ancestors.

For each protein pair, we take the maximum score between the two matrices (MICA sores and NSP scores) to be the co-membership score of the protein pair in the co-membership clique. For two proteins *p*1 and *p*2 annotated by pathways *P*1 and *P*2, the co-membership score of *p*1 and *p*2 is equal to:

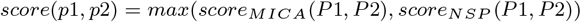

In the end, co-membership interactions combine MICA and NSP scores. A score of zero between two proteins indicates that they are only members of the root pathway without any `abstraction:NextStepPathway` relationship.

### Benchmarking pathway co-membership methods

In order to benchmark pathway co-membership methods and evaluate our approach, we conducted gene sets prioritization experiments using ten cancer hallmarks gene sets, that were downloaded from https://cancerhallmarks.com/. The ten cancer hallmarks gene sets are : “Tissue invasion and metastasis”, “Reprogramming energy metabolism”, “Replicative immortality”, “Genome instability”, “Resisting cell death”, “Evading growth suppressors”, “Evading immune destruction”, “Sustained angiogenesis”, “Sustained proliferative signaling” and “Tumor promoting inflammation”. Genes in each of the ten cancer hallmarks gene sets were converted to UniProt identifiers using the UniProt ID mapping tool (Zaru et al., 2023) on the UniProt knowledge base website^4^.

We chose the random walk with restart (RWR) method to conduct gene prioritization experiments. For the RWR experiments and for each cancer hallmark gene set, 80% of the genes were used as seeds and 20% were used as targets. RWR were run on five different pathway co-membership cliques of Reactome (version 96, April 2026):

- The Reactome pathway co-membership clique scored with our approach taking into account hierarchical and sequential information of pathways
- The Reactome pathway co-membership clique scored with a variation of our scoring approach taking into account only the hierarchical information of pathways
- The Reactome pathway co-membership clique scored with a Cosine similarity measure
- The Reactome pathway co-membership clique scored with a Jaccard similarity measure
- The unweighted Reactome pathway co-membership clique

We used the MultiXrank Python package (Baptista et al., 2022) to run the RWR. The global restart probability was set to 0.1 on order to allow a long-range exploration of the networks. The same experiment was also conducted with a restart probability of 0.7 in order to asses the performances of the metrics for a close neighborhood exploration task (results shown in Supplementary Figure S-2). RWR results were analyzed with enrichment curves that show how much targets genes were highly scored by the RWR (by calculating areas under the curves).

## Results

### Respective influence of compositional and sequential information: “Signaling by TGF-beta receptor complex” pathway from Reactome as a use case

To illustrate our weighted co-membership approach, we detail in this section the construction of the weighted co-membership clique of the “Signaling by TGF-beta receptor complex” pathway from Reactome (R-HSA-170834). This pathway is composed of six sub-pathways, and annotates a total of 93 proteins (UniProt identifiers counted in the BioPAX file).

We constructed the pathway-centered abstraction of the pathway “Signaling by TGF-beta receptor complex”, following the methods detailed in the previous sections. The Cytoscape (Shannon et al., 2003) visualization of the pathway-centered abstraction is presented in Figure 3, panel A. On this figure, the “Signaling by TGF-beta receptor complex” pathway is represented on top of the graph since it is the root pathway. The six sub-pathways are laid out hierarchically below the root pathway. The size of the pathways depends on the number of proteins they annotate. Their color (gradient of green) depends on their *score*_*MICA*_ (lower when green is light and higher when green is dark). Since the *score*_*MICA*_ of pathways depends on their specificity in the proteins they annotate, we retrieve higher *score*_*MICA*_ values associated to leaf pathways and lower values associated to pathways higher in the hierarchy (the root pathway has a *score*_*MICA*_ value of zero). The composition edges of the abstraction (`abstraction:IsAComponentOf` relationships) are depicted with gray diamond arrow. Their width and transparency vary according to their weight computed with the information content in proteins formula defined in the previous sections. Pathways can also be linked by `abstraction:NextStepPathway` edges (depicted in red on the figure, their width and transparency vary according to their weight).

Figure 3 panel B presents the distribution of scores in the obtained weighted co-membership clique. The provenance of scores is also detailed. Green bars correspond to co-membership scores from the MICA. Red bars correspond to co-membership scores from the `abstraction:NextStepPathway` edges (NSP scores). The maximal weights correspond to pairs of proteins that are as close as possible in the clique (i.e proteins that are annotated by the same subpathway). These weights are thus always originating from the score of the MICA (which is the parent pathway of the two proteins). In the clique of the “Signaling by TGF-beta receptor complex” pathway, these maximal scores are found between 0.4 and 0.76 and are represented by short green bars on the right of the histogram. The maximal value of co-membership score is found for proteins belonging to the pathway “TGF-beta receptor signaling in EMT (epithelial to mesenchymal transition)” that is the leaf pathway with the highest *score*_*MICA*_. On the contrary, the minimal value of co-membership scores (zero scores) is found for proteins only linked by the root pathway (i.e no `abstraction:NextStepPathway` edge linking parent pathways, and no MICA except root pathway).

Also, we retrieve co-membership scores corresponding to *score*_*NSP*_ values (red bars). The number of red bars corresponds to the number of different NSP scores fount the in the pathway-centered abstraction of Figure 2. Some co-membership scores can have multiple provenances: it is possible that the same score can be assigned in a specific situation between two proteins because of the presence of a `abstraction:NextStepPathway` edge between pathways, or in another situation with another protein pair because scores of the presence of a MICA score having the same value. In our example, this situation is represented by bi-colored bars (at the scores values of 0.24 and 0.26). Also, in the case of two proteins annotated by two pathways sharing a `abstraction:NextStepPathway` edge and having a direct MICA, the co-membership score can come either from the *score*_*MICA*_ or the *score*_*NSP*_ .

**Figure 2.**
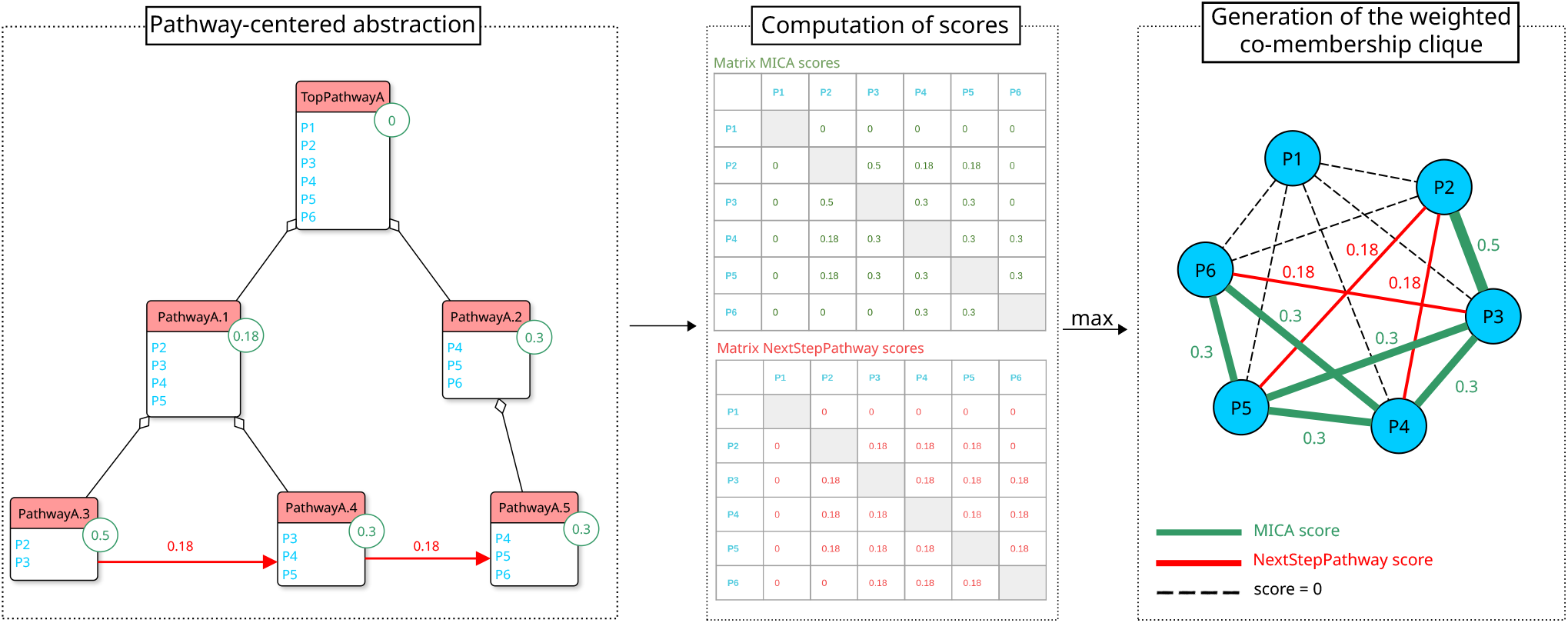
Methods used to generate a weighted co-membership clique from a pathway. The abstracted representation of a pathway toy example is represented in the left panel of the figure. Individual pathways are depicted using boxes with a read header containing their label. The proteins associated to each pathway are listed inside the boxes. Hierarchical relationships between pathways (`abstraction:IsAComponentOf edges`) are represented using black diamond arrows. Sequential relationships between pathways (`abstraction:NextStepPathway edges`) are represented using red arrows. On each pathway, the score in information content of proteins (*score*_*MICA*_) is mentioned in a green tag. Scores of the `abstraction:NextStepPathway edges` (*score*_*NSP*_ ) are mentioned in red above the corresponding arrows. In the middle panel of the figure are presented the matrices used to compute the scores between pairs of proteins in the pathway. Pairwise scores between proteins are computed using MICA scores and NSP scores, and the maximum of both scores is taken as a weight in the final weighted co-membership clique presented in the right panel of the figure. Co-membership edges weighted with a MICA score are depicted in green and those scored with a NSP score are depicted in red. Co-membership interactions with a score of zero are depicted as dashed black edges. MICA = Most Informative Common Ancestor, NSP = NextStepPathway

**Figure 3.**
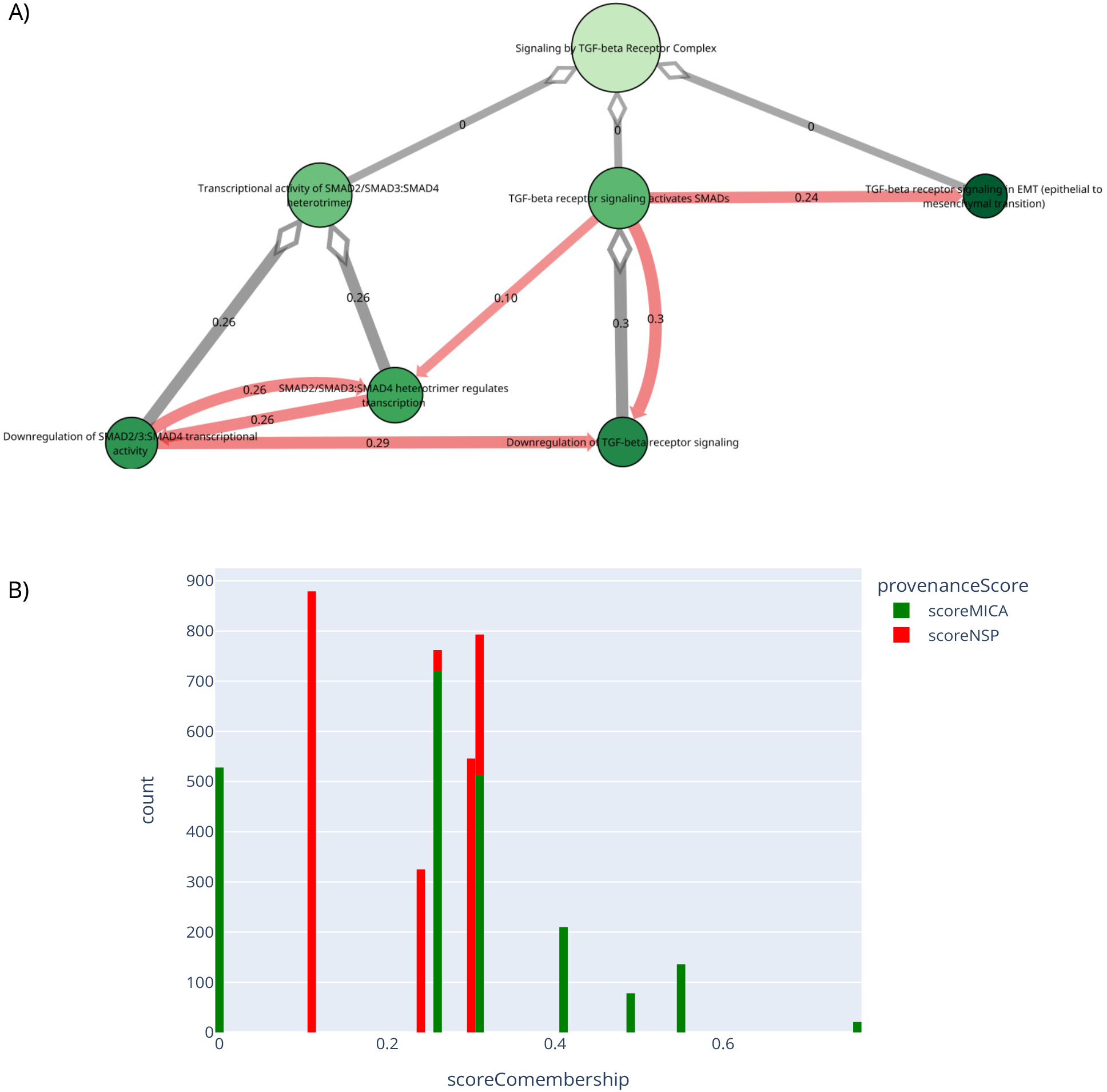
**Panel A: Pathway-centered abstraction of the pathway “Signaling by TGF-beta Receptor Complex” from Reactome** `R-HSA-170834`. Pathways are represented as nodes, for which the size depends on the number of proteins annotated by this pathway. The color of the nodes reflects the MICA score of the pathway (lighter green = low MICA score, darker green = higher MICA score). `abstraction:IsAComponentOf` edges are represented using grey diamond arrows, and `abstraction:NextStepPathway` edges are represented using red arrows. The corresponding weights are labeled on the edges, and are also reflected by the transparency of the edges. Visualization obtained with Cytoscape. **Panel B: Distribution of co-membership scores for the pathway “Signaling by TGF-beta receptor complex” from Reactome (R-HSA-170834)**. The bars represent the proportion of scores found in the weighted co-membership clique of the pathway “Signaling by TGF-beta receptor complex”. Green bars correspond to scores originating from MICA scores and red bars correspond to scores originating from `abstraction:NextStepPathway` (NSP) scores.

We then constructed the co-membership clique of the “Signaling by TGF-beta receptor complex” pathway and obtain a clique of 4, 278 weighted interactions between the 93 proteins. We clustered the obtained protein pairs using their co-membership score (cf Figure 4). The clustermap presents the 93 proteins of the “Signaling by TGF-beta receptor complex” pathway clustered by pairs. The heatmap is colored depending the co-membership score of the protein pairs. The pathways annotating each protein is also represented using the grid on the top of the clustermap: a black cell in the grid means that the corresponding proteins belongs to the line of the pathway. The “Signaling by TGF-beta receptor complex” pathway contains all of the 93 proteins so we observe a black line on the grid. We observe that the maximum co-membership scores, represented as the yellow cluster on the figure, are found for pairs of proteins located in the pathway “TGF-beta receptor signaling in EMT (epithelial to mesenchymal transition)”, confirming our observations on Figure 3 A (higher MICA scores). The three green clusters correspond to protein pairs located in the other leaf pathways (“Downregulation of TGF-beta receptor signaling”, “SMAD2/SMAD3:SMAD4 heterotrimer regulates transcription” and “Downregulation of SMAD2/3:SMAD4 transcriptional activity”). Details about proteins composing each clusters are provided in Supplementary File S-1.

**Figure 4.**
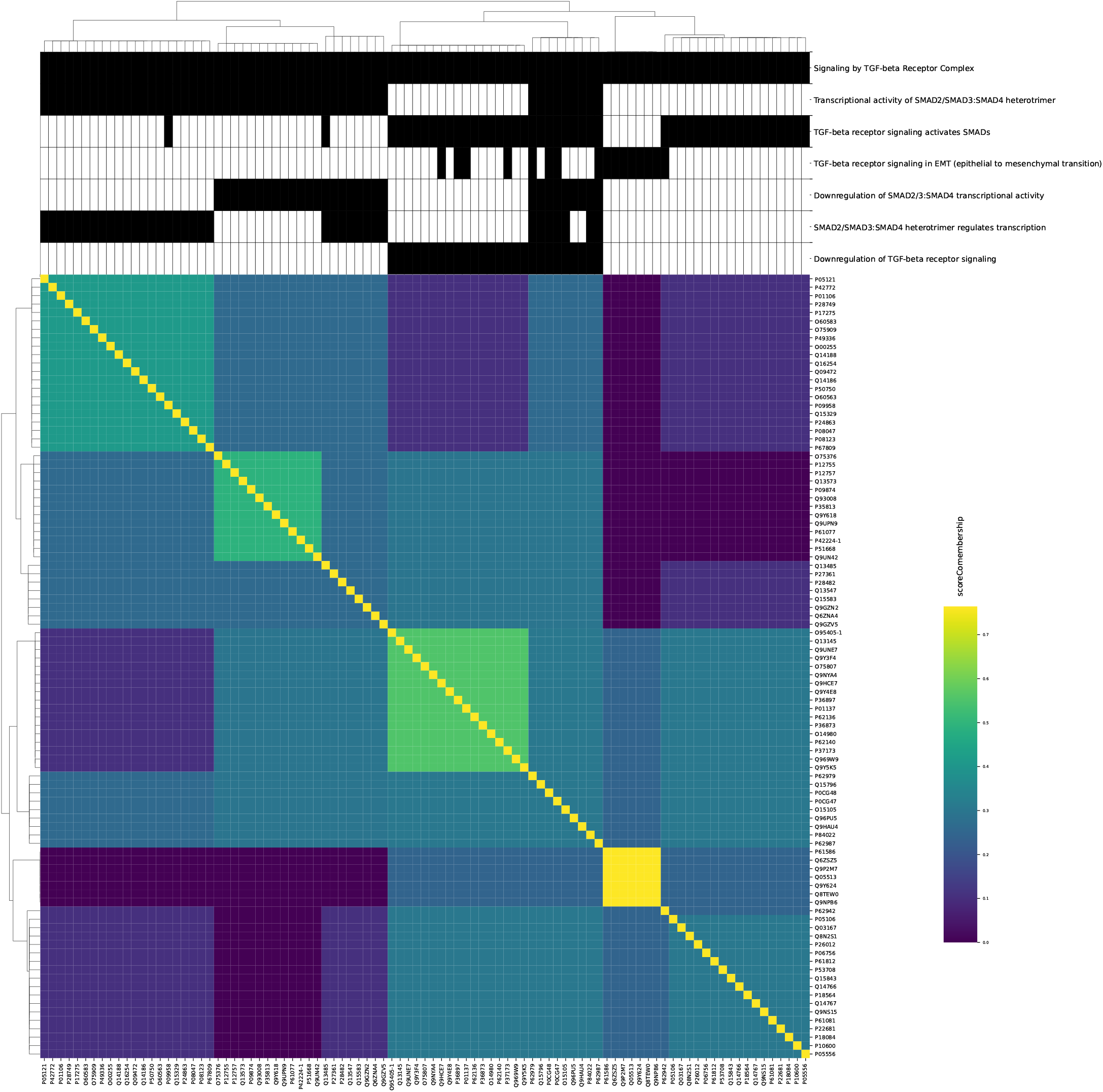
Clustermap of co-membership scores for the pathway “Signaling by TGF-beta receptor complex” from Reactome (R-HSA-170834). The 93 proteins of the “Signaling by TGF-beta receptor complex” pathway are clustered in rows and columns depending on their pairwise co-membership score. Cells of the clustermap are colored with regard to the co-membership score following a color gradient depicted on the right of the figure. For each protein, the most specific pathway in the hierarchy associated with this protein is depicted with a black cell in the grids on the top of the figure.

### Generalization: scoring co-membership interactions of the 29 Reactome top-level pathways

In this section, we detail how we used our scoring system to generate the weighted pathway co-membership cliques of the 29 top-level pathways of Reactome. Table 1 presents some metrics about the pathway co-membership cliques of the 29 Reactome top-level pathways. The number of proteins annotated by each top-level pathway is detailed, as well as the size of the generated cliques, and the number of scores originating from a MICA score or a NSP score. We remark a significant heterogeneity, both in the sizes of the generated cliques, and in the distribution of MICA and NSP scores. The top-level pathways “Circadian Clock” and “DNA Replication” show the highest NSP scores (approximately 40%, 38.4% and 36% respectively), reflecting that their sub-pathways are well interconnected. On the contrary, the top-level pathways “Immune System” presents a very few amount of NSP scores (2.2%), and there are no NSP scores at all for “Drug ADME” and “Portein Localization” pathways. These pathways present very few or no `bp3:nextStep` relationships.

**Table 1.** Metrics about the pathway co-membership cliques of the 29 top-level pathways of Reactome (version 96). The number of proteins (UniProt identifiers) associated to each top-level pathway is detailed in the first column. The size of the co-membership cliques of each top-level pathway is detailed in the second column. In the two last columns, the number of MICA scores and NSP scores in the co-membership cliques are detailed.

| Top-pathway | Reactome ID | Nb of proteins | Clique size | Nb MICA scores | Nb NSP scores | % of NSP scores |
| --- | --- | --- | --- | --- | --- | --- |
| Autophagy | R-HSA-9612973 | 140 | 9,730 | 9,021 | 709 | 7.3% |
| Cell Cycle | R-HSA-1640170 | 665 | 220,780 | 186,355 | 34,425 | 15.6% |
| Cell-Cell Communication | R-HSA-1500931 | 322 | 51,681 | 47,684 | 3,997 | 7.7% |
| Cellular Response To Stimuli | R-HSA-8953897 | 868 | 376,278 | 360,039 | 16,239 | 4.3% |
| Chromatin Organization | R-HSA-4839726 | 345 | 59,340 | 58,958 | 382 | 0.64% |
| Circadian Clock | R-HSA-9909396 | 111 | 6,105 | 3,763 | 2,342 | 38.4% |
| Developmental Biology | R-HSA-1266738 | 1,416 | 1,001,820 | 965,231 | 36,589 | 3.6% |
| Digestion And Absorption | R-HSA-8963743 | 28 | 378 | 327 | 51 | 13.5% |
| Disease | R-HSA-1643685 | 2,533 | 3,206,778 | 3,167,004 | 39,774 | 1.24% |
| DNA Repair | R-HSA-73894 | 314 | 49,141 | 41,696 | 7,445 | 15.15% |
| DNA Replication | R-HSA-69306 | 153 | 11,628 | 7,434 | 4,194 | 36% |
| Drug ADME | R-HSA-9748784 | 111 | 6,105 | 6,105 | 0 | 0% |
| Extracellular Matrix Organization | R-HSA-1474244 | 310 | 47,895 | 47,328 | 567 | 1.18% |
| Gene Expression(Transcription) | R-HSA-74160 | 1,537 | 1,180,416 | 1,154,481 | 25,935 | 2.2% |
| Hemostasis | R-HSA-109582 | 717 | 256,686 | 224,015 | 32,671 | 12.7% |
| Immune System | R-HSA-168256 | 2,311 | 2,669,205 | 2,609,520 | 59,685 | 2.2% |
| Metabolism | R-HSA-1430728 | 2,216 | 2,454,220 | 2,386,699 | 67,521 | 2.7% |
| Metabolism Of Proteins | R-HSA-392499 | 2,113 | 2,231,328 | 2,188,362 | 42,966 | 1.9% |
| Metabolism Of RNA | R-HSA-8953854 | 747 | 278,631 | 251,066 | 27,565 | 10% |
| Muscle Contraction | R-HSA-397014 | 198 | 19,503 | 17,371 | 2,132 | 11% |
| Neuronal System | R-HSA-112316 | 400 | 79,800 | 77,592 | 2,208 | 2.8% |
| Organelle Biogenesis And Maintenance | R-HSA-1852241 | 285 | 40,470 | 39,901 | 569 | 1.4% |
| Programmed Cell Death | R-HSA-5357801 | 203 | 20,503 | 17,871 | 2,632 | 12.8% |
| Protein Localization | R-HSA-9609507 | 166 | 13,695 | 13,695 | 0 | 0% |
| Reproduction | R-HSA-1474165 | 127 | 8,001 | 7,935 | 66 | 0.8% |
| Sensory Perception | R-HSA-9709957 | 639 | 203,841 | 202,746 | 1,095 | 0.5% |
| Signal Transduction | R-HSA-162582 | 2,586 | 3,342,405 | 3,145,924 | 196,481 | 5.9% |
| Transport Of Small Molecules | R-HSA-382551 | 716 | 255,970 | 251,602 | 4,368 | 1.7% |
| Vesicle-Mediated Transport | R-HSA-5653656 | 730 | 266,085 | 262,702 | 3,383 | 1.3% |

### Scaling the weighted co-membership system to the entire Reactome knowledge base

In order to compute the weighted pathway co-membership clique of the entire Reactome pathway knowledge base, we worked on the entire BioPAX model. We applied the scoring system described in the Methods section, but we changed the total number of proteins in the denominator of the information content in proteins formula. Instead of dividing by the total number of proteins in the pathway, we divide by the total number of proteins in Reactome (12102 UniProt identifiers counted in the version 96 of Reactome). Moreover, since we work on the entire Reactome knowledge base, the 29 Reactome top-level pathways get interconnected by additional `abstraction:NextStepPathway` edges that are not described when working on individual top-level pathways. We thus extracted those inter-top pathways NSP scores (that we names NSP inter-top). For more detailed information on this scaling process, an explicative diagram is provided as Supplementary Figure S-3.

The global co-membership clique of Reactome (version 96) contains 73, 223, 151 weighted interactions. Among those, 15, 353, 715 are interactions scored by a MICA score (20%) and 132, 634 interactions are scored by a NSP score (0.2%). Also, it is interesting to note that 277, 701 interactions have a score originating from a NSP inter-top pathways (0.4%), meaning that these specific protein interactions have a score enriched through the presence of sequences between their associated top-level pathways. The rest of the co-membership interactions in the clique (57, 459, 101 interactions) are scored with a zero score (78%).

### Benchmark of gene prioritization based on pathway co-membership

In this section we evaluate our weighted pathway co-membership. We aim at demonstrating that groups of proteins supposed to be biologically close are closer than by chance in the weighted co-membership clique of Reactome. To do this, we used ten cancer hallmarks gene sets as genes to be prioritized, and used a random walk with restart (RWR) approach as a gene prioritization method. The cancer hallmarks are: “Tissue invasion and metastasis”, “Reprogramming energy metabolism”, “Replicative immortality”, “Genome instability”, “Resisting cell death”, “Evading growth suppressors”, “Evading immune destruction”, “Sustained angiogenesis”, “Sustained proliferative signaling” and “Tumor promoting inflammation”.

We evaluated gene prioritization performance by applying Random Walks with Restart (RWR) to weighted and unweighted versions of the Reactome pathway co-membership cliques. Using four distinct weighting methods, we tested two scenarios: a low restart probability (*r* = 0.1) to assess long-range network exploration (Figure 5), and a high restart probability (*r* = 0.7) for local neighborhood detection (Supplementary Figure S-2). Cancer hallmarks served as seeds (20%) and targets (80%).

**Figure 5.**
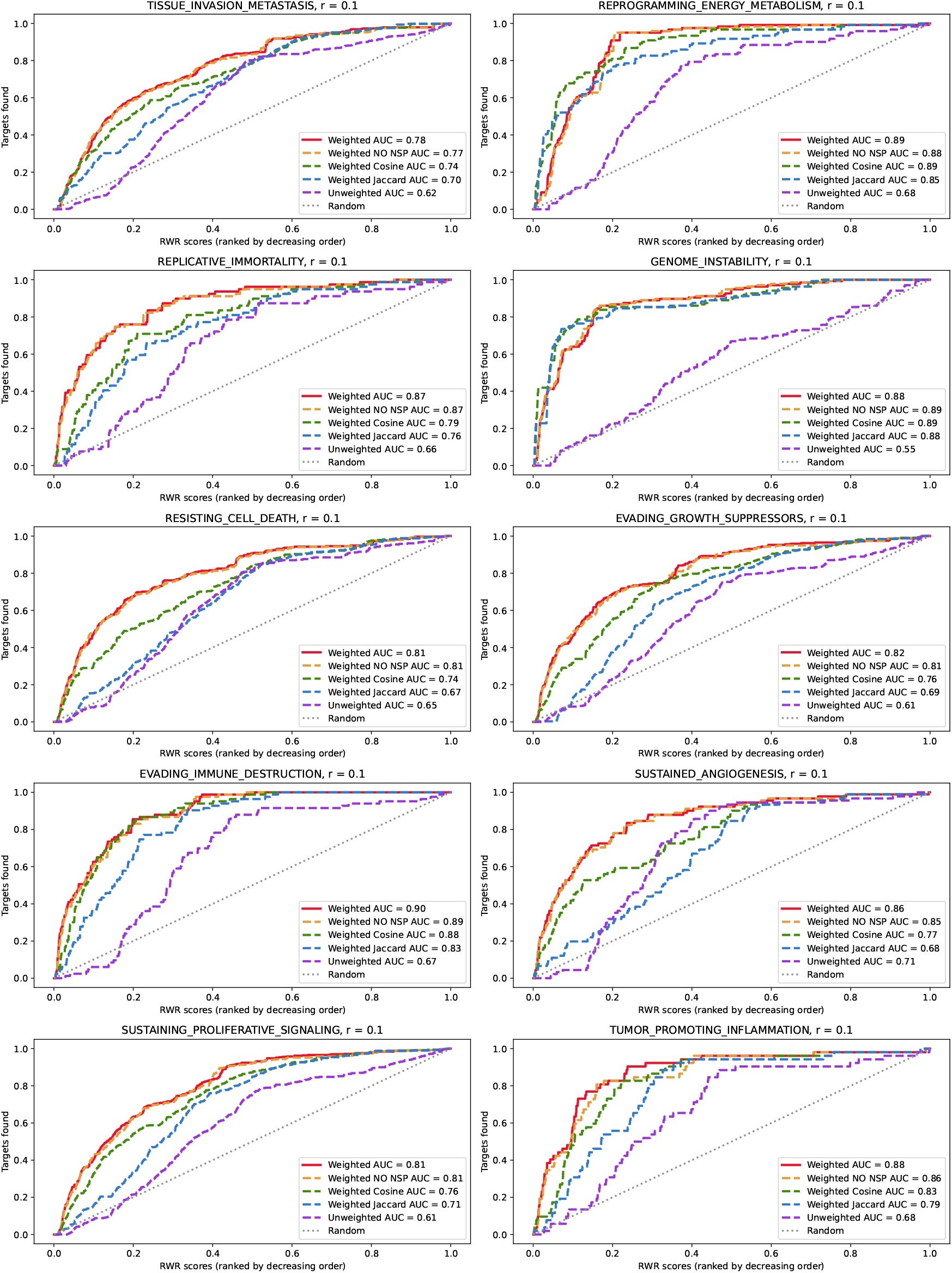
Enrichment curves of the random walk with restart scores for different scoring methods of the co-membership clique of Reactome (version 96). Nine cancer hallmarks gene sets were used as seeds (20%) and targets (80%) for the RWR (“Tissue invasion and metastasis”, “Reprogramming energy metabolism”, “Replicative immortality”, “Genome instability”, “Resisting cell death”, “Evading growth suppressors”, “Evading immune destruction”, “Sustained angiogenesis”, “Sustained proliferative signaling” and “Tumor promoting inflammation”). The four co-membership interactions scoring methods are: our scoring system based on the hierarchical information of Reactome pathways and their sequentiality (plain red curve), the same scoring system without taking into account the sequentiality of pathways (dashed orange curve), the Cosine similarity (dashed green curve) and the Jaccard similarity (dashed blue curve). The dashed purple curve represents the RWR results obtained with the unweighted co-membership clique of Reactome. The dotted gray line represents random results.

Figure 5 presents the enrichment curves obtained with the RWR results for the ten hallmarks and on the five Reactome pathway co-membership cliques: : the clique scored with our approach (plain red curve), the clique scored with our approach without taking into account the sequentiality of pathways (dashed orange curve), the clique scored with a Cosine similarity (dashed green curve), the clique scored with a Jaccard similarity (dashed blue curve) and the unweighted clique (dashed purple curve). The gray dotted line represents the random situation where target hallmarks are randomly arranged in the results of the scores. The RWR scores are sorted in descending order on the x-axis. The y-axis represents the cumulative count of target hallmark genes recovered, and the curves steps up at each position corresponding to a score matched by a target gene, with the step height proportional to the number of targets sharing that score. For each hallmark and each Reactome clique, we computed the area under the curve (AUC).

Globally, the better AUC is always obtained with our approach (plain red curves) for the ten cancer hallmarks gene sets prioritized (AUC always found between 0.78 (for “Tissue Invasion Metastasis”) and 0.90 (for “Evading Immune Destruction”)). This reflects the benefits of taking into account both the hierarchy of pathways and their sequence in the co-membership scoring system. When modifying our approach to only take into account the hierarchy and not the sequences (dashed orange curves), the performances stays equivalent or slightly decrease (AUC between 0.77 (for “Tissue Invasion Metastasis”) and 0.89 (for “Genome Instability”)). These results reflect the fact that NSP scores represent a few amount of the total scores in the Reactome weighted clique (0.4%), so their contribution to the scores is low. However, taking NSP scores into account still allows to slightly increase the results. RWR results on the co-membership cliques scores with state-of-the art methods Cosine and Jaccard similarity present lower performances than our method for this task (respectively dashed green and blue curves). For the Cosine similarity, AUC are comprised between 0.74 (for “Tissue Invasion Metastasis” and “Resisting Cell Death”) and 0.89 (for “Reprogramming Energy Metabolism” and “Genome Instability”). For the Jaccard similarity, AUC are comprised between 0.67 (for “Resisting Cell Death”) and 0.88 (for “Genome Instability”). RWR results on the unweighted co-membership clique of Reactome show performances close to random (from 0.55 (for “Genome Instability”) to 0.71 (for “Sustained Angiogenesis”). It shows that not weighting co-membership interactions introduce a lacking of distance between proteins.

## Discussion

### Pathway sequences and composition are highly informative

Performing gene prioritization on the weighted pathway-co-membership cliques of Reactome scored with our approach allowed to evaluate the benefits of taking into account both the hierarchy and the sequences of pathways for scoring co-membership interactions. By showing a greater AUC for all cancer hallmarks gene sets, our method based on the hierarchy and sequences of pathways tends to be efficient for capturing biological distance between proteins in the co-membership clique. When we applied a modified version of our method that only considered MICA scores, we observed a slight loss of performance. At the scale of the Reactome pathway co-membership clique, NSP scores only represent 0.6% of the scores (including NSP scores and NSP inter top-level pathways scores). Their contribution to the scores remain low, but since we still observe a slight increase of performances when adding them, we can consider that they bring significant information. Also, integrating NSP inter top-level pathway scores makes 0.4% of the interactions to pass from a zero-score to a non-null score.

In Reactome top-level pathways, knowledge structure can strongly vary, either by prioritizing compositional relationships, sequential relationships, or both. For example, in the “Circadian Clock” top-level pathway (`R-HSA-9909396`), we have pathway with very few but strongly sequential components (9 sub-pathways and 111 proteins). On the contrary, other top-level pathways like “Reproduction” (`R-HSA-1474165`) present very few sequential edges and are strongly composed (8 sub-pathways and 127 proteins). For the pathways using a lot sequential relationships, taking them into account in the scoring system can have a strong influence on co-membership weights. Differences in representation may be linked to different biological realities that should not be distorted in order to improve the metric. An efficient co-membership metric should take into account both the composition and the hierarchy of pathways.

### Benefits of our approach relatively to state-of-the art methods

Jaccard and Cosine similarity measures show lower performance than our method on such a long-range network exploration task. However, when performing the same gene prioritization experiment using a restart probability of 0.7, we observe that all weighting methods show equivalent performances (see Supplementary Figure S-2). Using a higher restart probability makes the random walk stay in the close neighborhood of the seeds, and is more suited for community detection tasks than for long-range network exploration. In this case, since all weighting metrics show equivalent performances, we suppose that all metrics tend to approximate well the distance between proteins and their close neighbors. However, when allowing the random walk to diffuse further through the networks with a restart probability of 0.1 (Figure 5), our method tends to be more efficient (higher AUC for all the ten hallmarks gene sets). Therefore, for close protein neighborhood exploration, our method is equivalent to state-of-the-art metrics, but outperforms them for long-range network exploration tasks.

### Choosing the co-membership scoring function

Another point that could be discussed is the choice of the co-membership scoring function. For scoring pathway co-membership interactions, we chose a function that takes the maximum score among the MICA scores and the potential NSP scores of the two protein’s parent pathways. Doing like this, we aim to reflect as much as possible the proximity of pathways and thus the proximity of associated proteins, either through their distance in the compositional hierarchy of pathways or through their interconnections. Other possible scoring functions could be summing the MICA and NSP scores, or doing a mean, rather than taking the maximum. We chose to take the maximum score in order to be independent from potential irregularities in the structure of pathways. We tested to weight co-membership interactions by taking the sum of MICA and NSP scores, and performed the same gene prioritization experiments but we obtained results equivalent to our method taking the maximum score (data not shown). However, these results could be specific to the Reactome BioPAX model and further work on other databases would be needed to assess if there are differences in both scoring functions.

### Perspectives

For getting some perspectives on our work, we can start by pointing the fact that generating pathway co-membership cliques can be a tedious task for complex and large pathways. For individual top-level pathways, the generation takes a few minutes, but when scaling to the entire Reactome weighted co-membership clique it can take few hours. However, the more intensive part of process is the generation of all protein pairs in a given pathway, and not the way they are scored. Still, the required computational resources can be cited as a limitation of our approach. Re-computing weighted pathway co-membership cliques is only necessary when the Reactome knowledge base is updated. Since Reactome is updated every three months, the task of automatically updating and re-computing pathway co-membership cliques is a serious question. For the moment, computing pathway co-membership cliques should be done manually and totally re-computed from scratch at every update. But this process could be automated in the future, and even simplified if strategies are found to avoid re-computing the score of every protein pair at each update.

Another perspective point on our approach is about its generalization to other pathway knowledge bases. Here we tested our pathway co-membership scoring system on Reactome pathways, and we showed that it works well on this knowledge base. However, since our approach is based on BioPAX which is a standard format adopted by the majority of pathway knowledge bases, we could imagine applying our co-membership scoring approach to other pathway resources. Still, it is important to note that not all pathway knowledge bases make as much use of the richness of BioPAX as Reactome, as pointed out in our article on the subject (Beust et al., 2024). Some pathway knowledge bases export their pathways in BioPAX, but lack richness either the description of their composition, or the description of their sequences. These incomplete pathway descriptions in BioPAX could hinder the use of our metric, but it stays generic and applicable to any pathway resource using BioPAX.

Finally, our pathway co-membership scoring method can also be used jointly with other types of biological information to gain insights into biological processes in a systematic way. We could imagine multiplexing weighted pathway co-membership cliques with other resources or other contextual abstractions of Reactome pathways in a multi-scale perspective.

## Conclusion

As a conclusion, we developed a scoring system of pathway co-membership cliques based on the hierarchical and sequential information of Reactome pathways in BioPAX. Our method allows to generate a weighted clique of co-membership interactions among proteins in a pathway. We showed that our approach outperforms state-of-the-art pathway co-membership methods in gene prioritization tasks. Also, we showed that taking into account sequentiality of pathways in addition to its composition adds a significant amount of information for scoring co-membership interactions.

## Abbreviations

BioPAX: Biological Pathway Exchange
MICA: Most Informative Common Ancestor
IC: Information Content
NSP: NextStepPathway

## Supplementary Material

- Supplementary Figure S-1: Illustration of the pathway co-membership scoring system requirements
- Supplementary Figure S-2: Gene prioritization experiment results using a restart probability of 0.7
- Supplementary Figure S-3: Schema of the scaling process of the weighted co-membership to the entire Reactome knowledge base
- Supplementary File S-1: Details about the clusters of proteins of Figure 5. The file can be downloaded at: https://github.com/CecileBeust/AbstractionReactomeTopPathways/blob/main/Results/nature.com/articles/s42005-022-00937-9.PathwayComembership/07_TGFbeta/SuppFileS-1.xlsx

## Data availability

The codes to generate the weighted pathway co-membership cliques of Reactome pathways are publicly available on GitHub^5^. The pathway co-membership cliques of the 29 Reactome top-level pathways are publicly accessible and downloadable on the NDEx network platform^6^.

## Author contributions statement

C.B., O.D, N.T and E.B. conceived the experiment(s), C.B. conducted the experiment(s) and generated the results. C.B., O.D, N.T and E.B. analysed the results. C.B. wrote the manuscript. C.B., O.D, N.T and E.B. revised the manuscript.

## Generative AI Statement

During the preparation of this article, the authors used RAGaRenn (generative AI tool of the University of Rennes) with a Mistral Small

3.2 24B model to improve textual clarity through reformulation. After using this tool, the authors revised and edited the content as needed and take full responsibility for the content of the publication.

## Acknowledgments

We thank the GenOuest bioinformatics platform for providing the computing infrastructure.

---

**Cécile Beust** is a PhD student as University of Rennes. She works on the abstraction of biological knowledge graphs in BioPAX into contextual views.

**Olivier Dameron** is a professor at University of Rennes. He works on knowledge representation and bioinformatics, and develops methods based on ontologies for analyzing biomedical data.

**Nathalie Théret** is a Research Director at Inserm in Rennes. She works on the bioinformatics modeling of biological processes, mainly involved in chronic liver diseases.

**Emmanuelle Becker** is a professor at University of Rennes. She works on the integration of homogeneous and heterogeneous data and the extraction of knowledge from biological graphs.

**Supplementary Figure S-1:**
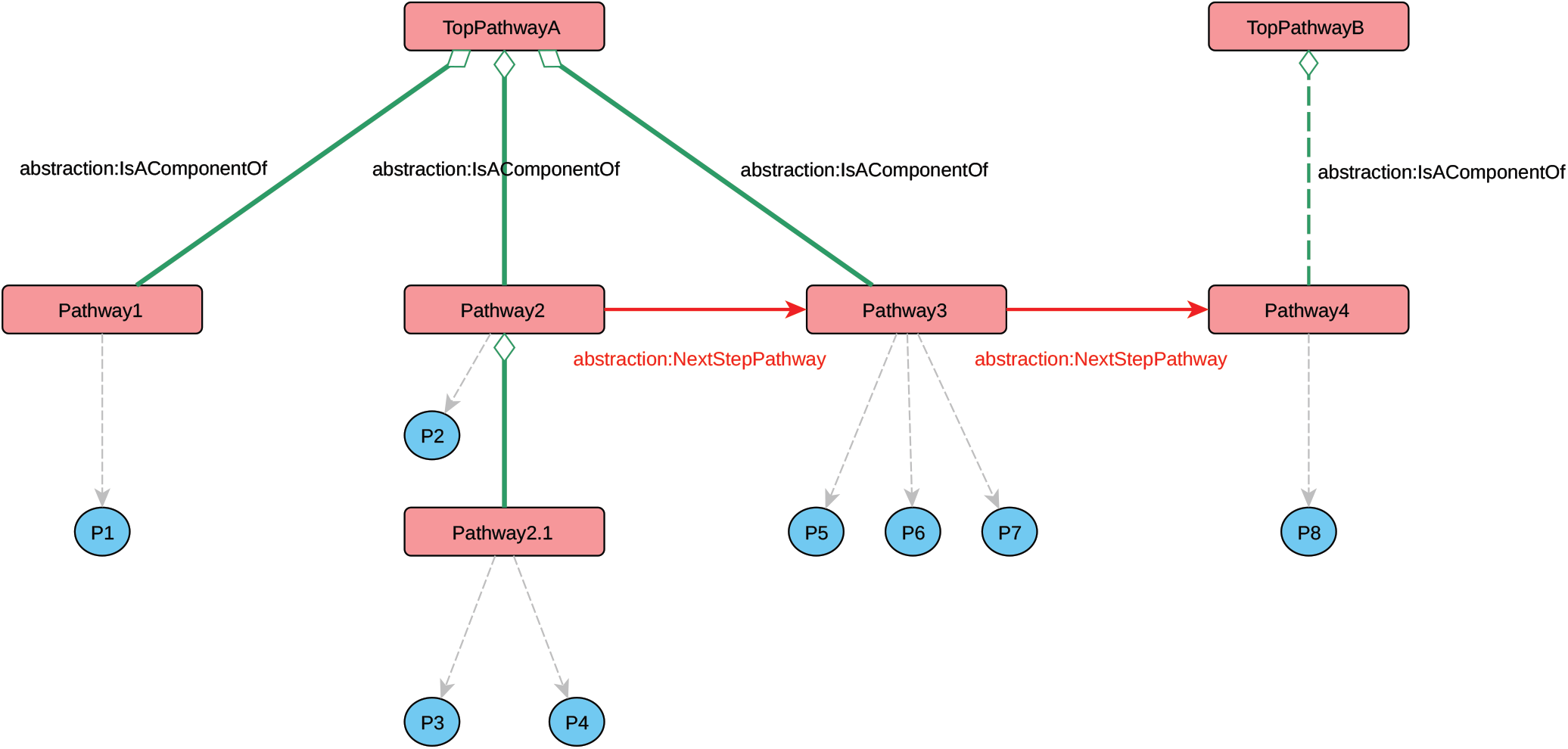
Illustration of the pathway co-membership scoring system requirements **Principle 1: Definition of the pathway co-membership** Principle 1: Definition of the pathway co-membership **Principle 2: Size of pathways matters** P3 and P4 share a stronger co-membership interaction than P6, P6 and P7 because the size of their associated pathway is smaller **Principle 3: Co-membership is a function of several scores** P2 and P4 participate in different pathways. Their co-membership score is a function of the scores of their associated pathways. **Principle 4: Sequentiality is taken into account** P3 and P4 are closer to P5 than P1 because of the presence of the abstraction:NextStepPathway edge connecting Pathway2 and Pathway3 P7 and P8 share a non-null co-membership interaction since Pathway3 and Pathway4 are connected by a *abstraction:NextStepPathway* edge

**Supplementary Figure S-2:**
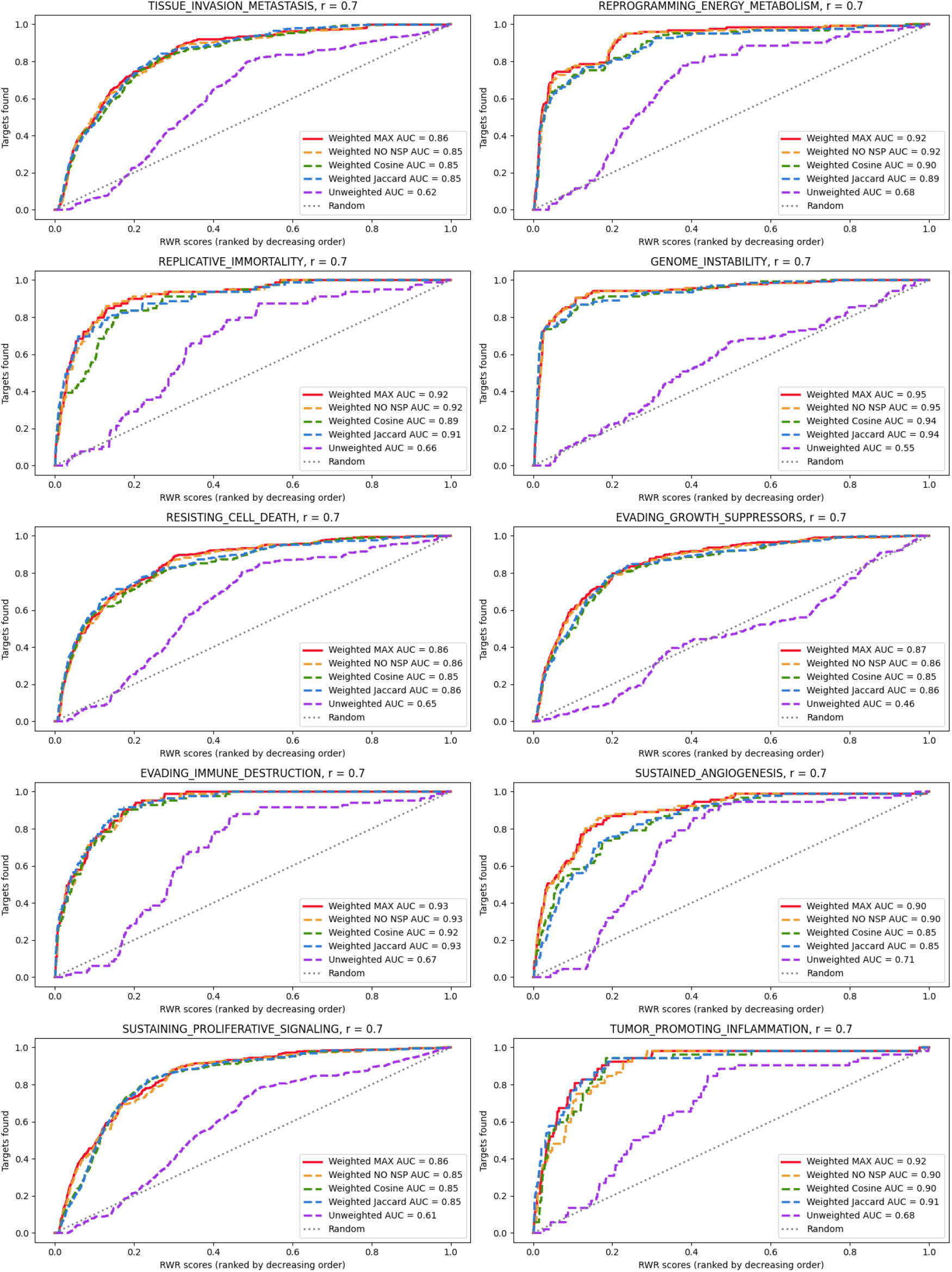
Gene prioritization experiment results using a restart probability of 0.7

**Supplementary Figure S-3:**
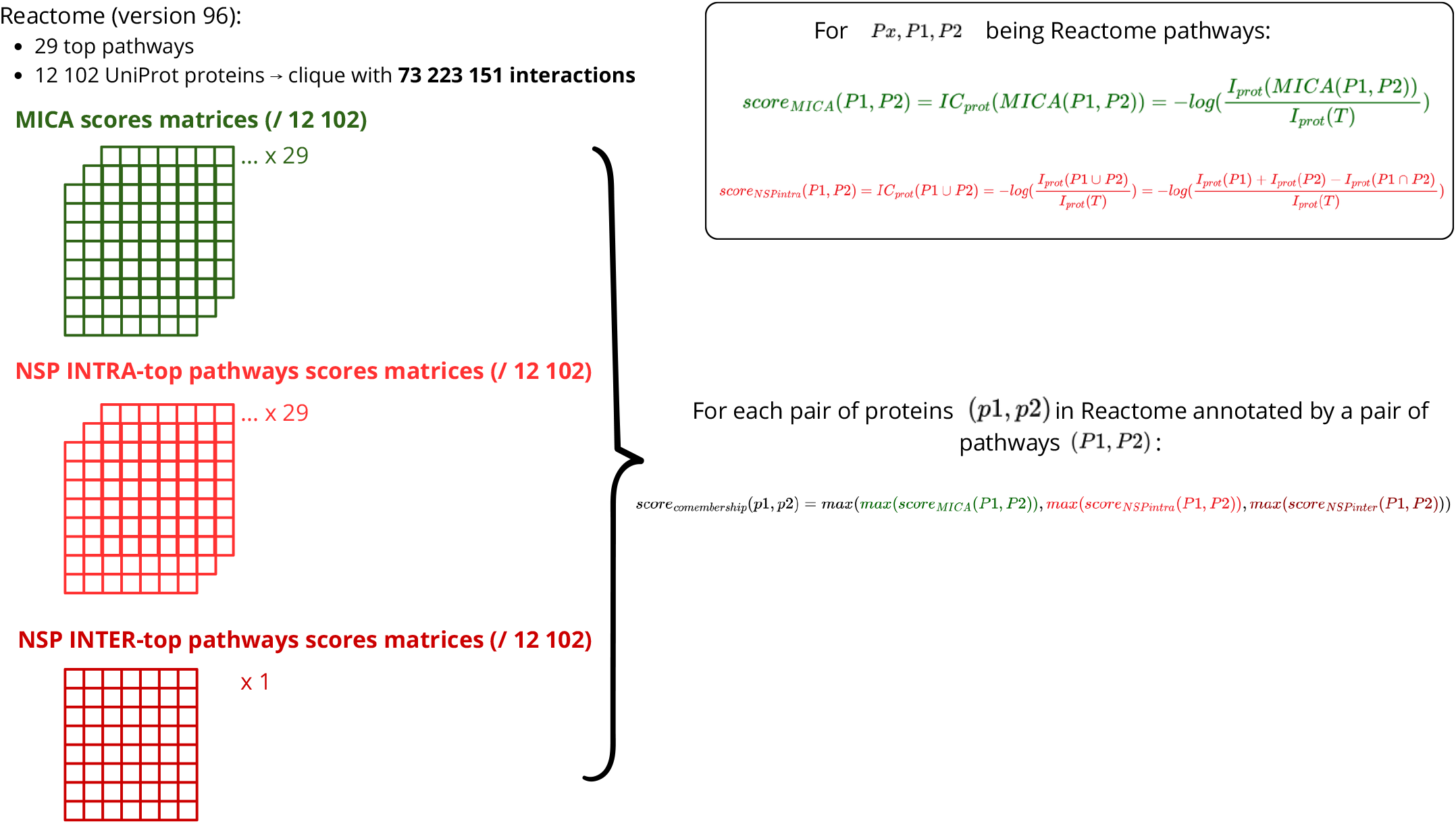
Schema of the scaling process of the weighted co-membership to the entire Reactome knowledge base

## Footnotes

* https://github.com/CecileBeust/AbstractionReactomeTopPathways.git

† https://www.ndexbio.org/folders/860c0b30-a222-11f1-94e3-005056ae3c32

‡ https://www.ndexbio.org/folders/860c0b30-a222-11f1-94e3-005056ae3c32

1 definition from the National Human Genome Research Institute: https://www.genome.gov/about-genomics/fact-sheets/Biological-Pathways-Fact-Sheet

2 https://download.reactome.org/96/biopax.zip

3 https://jena.apache.org/documentation/fuseki2/

4 https://www.uniprot.org/id-mapping

5 https://github.com/CecileBeust/AbstractionReactomeTopPathways.git

6 https://www.ndexbio.org/folders/860c0b30-a222-11f1-94e3-005056ae3c32

