## Supplementary Figure 1 for "Refining Protein Co-Membership in Reactome: A Weighted Scoring Approach Based on Hierarchy and Sequences of Patwhays"

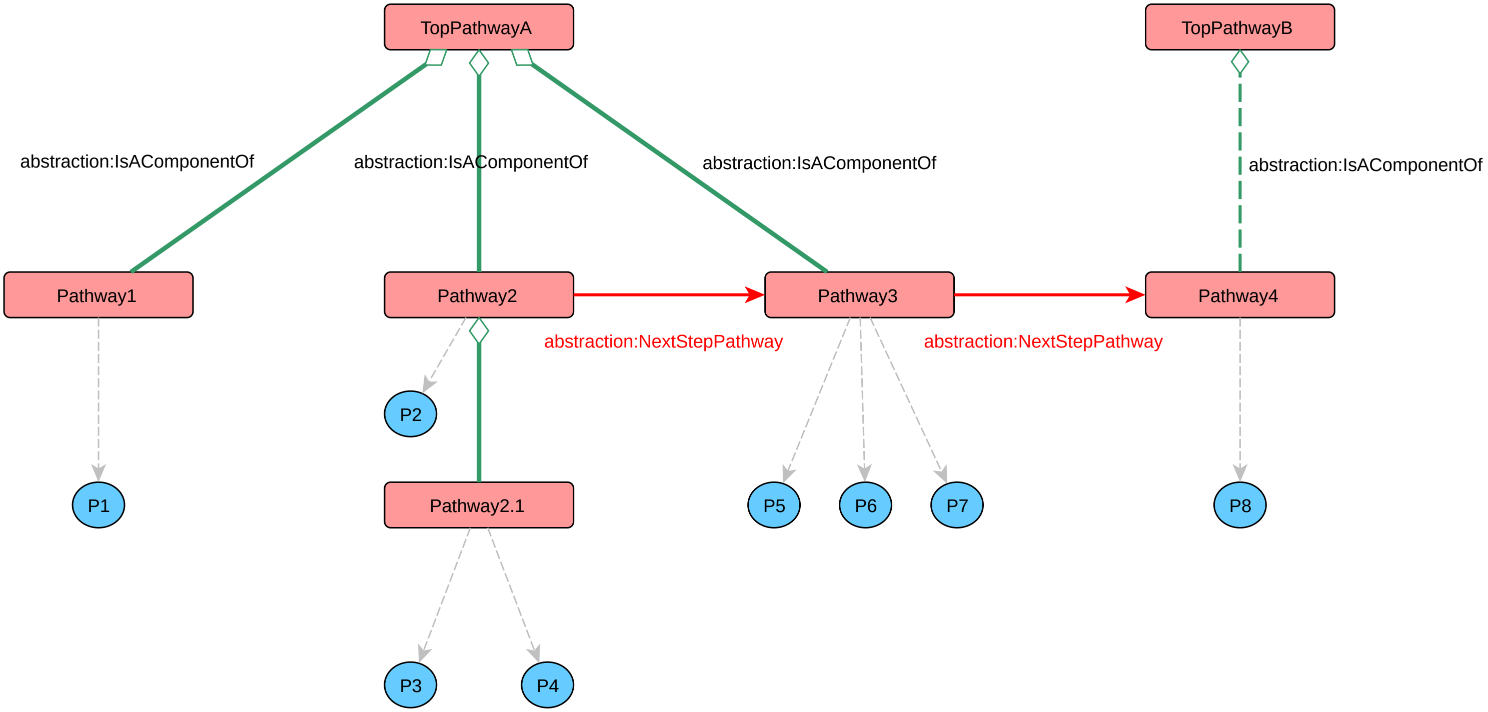


**Principle 1: Definition of the pathway co-membership**

- P3 and P4 are close because they are associated to the same pathway

**Principle 2: Size of pathways matters**

- P3 and P4 share a stronger co-membership interaction than P6, P6 and P7 because the size of their associated pathway is smaller

**Principle 3: Co-membership is a function of several scores**

- P2 and P4 participate in different pathways. Their co-membership score is a function of the scores of their associated pathways.

**Principle 4: Sequentiality is taken into account**

- P3 and P4 are closer to P5 than P1 because of the presence of the *abstraction:NextStepPathway* edge connecting Pathway2 and Pathway3

- P7 and P8 share a non-null co-membership interaction since Pathway3 and Pathway4 are connected by a *abstraction:NextStepPathway* edge
