## Supplementary Figure 3 for "Refining Protein Co-Membership in Reactome: A Weighted Scoring Approach Based on Hierarchy and Sequences of Patwhays"

Reactome (version 96):

- 29 top pathways
- 12 102 UniProt proteins → clique with **73 223 151 interactions**

### MICA scores matrices (/ 12 102)

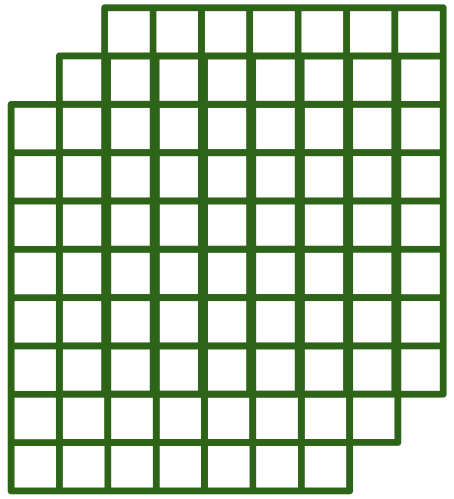

... x 29

### NSP INTRA-top pathways scores matrices (/ 12 102)

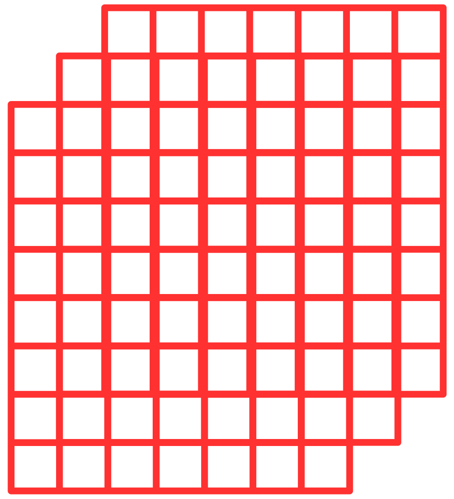

... x 29

### NSP INTER-top pathways scores matrices (/ 12 102)

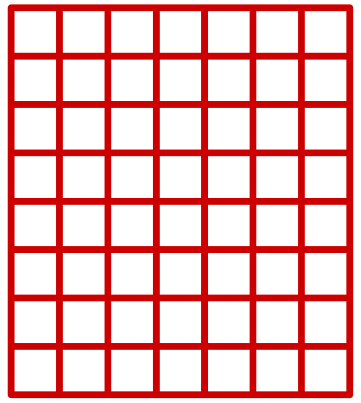

x 1

For $Px, P1, P2$ being Reactome pathways:

$$score_{MICA}(P1, P2) = IC_{prot}(MICA(P1, P2)) = -log(\frac{I_{prot}(MICA(P1, P2))}{I_{prot}(T)})$$

$$score_{NSPintra}(P1, P2) = IC_{prot}(P1 \cup P2) = -log(\frac{I_{prot}(P1 \cup P2)}{I_{prot}(T)}) = -log(\frac{I_{prot}(P1) + I_{prot}(P2) - I_{prot}(P1 \cap P2)}{I_{prot}(T)})$$

For each pair of proteins $(p1, p2)$ in Reactome annotated by a pair of pathways $(P1, P2)$:

$$score_{comembership}(p1, p2) = max(max(score_{MICA}(P1, P2)), max(score_{NSPintra}(P1, P2)), max(score_{NSPinter}(P1, P2)))$$
