## Supplementary figures and images for "Refining Protein Co-Membership in Reactome: A Weighted Scoring Approach Based on Hierarchy and Sequences of Patwhays"

### Supplementary Figure 2

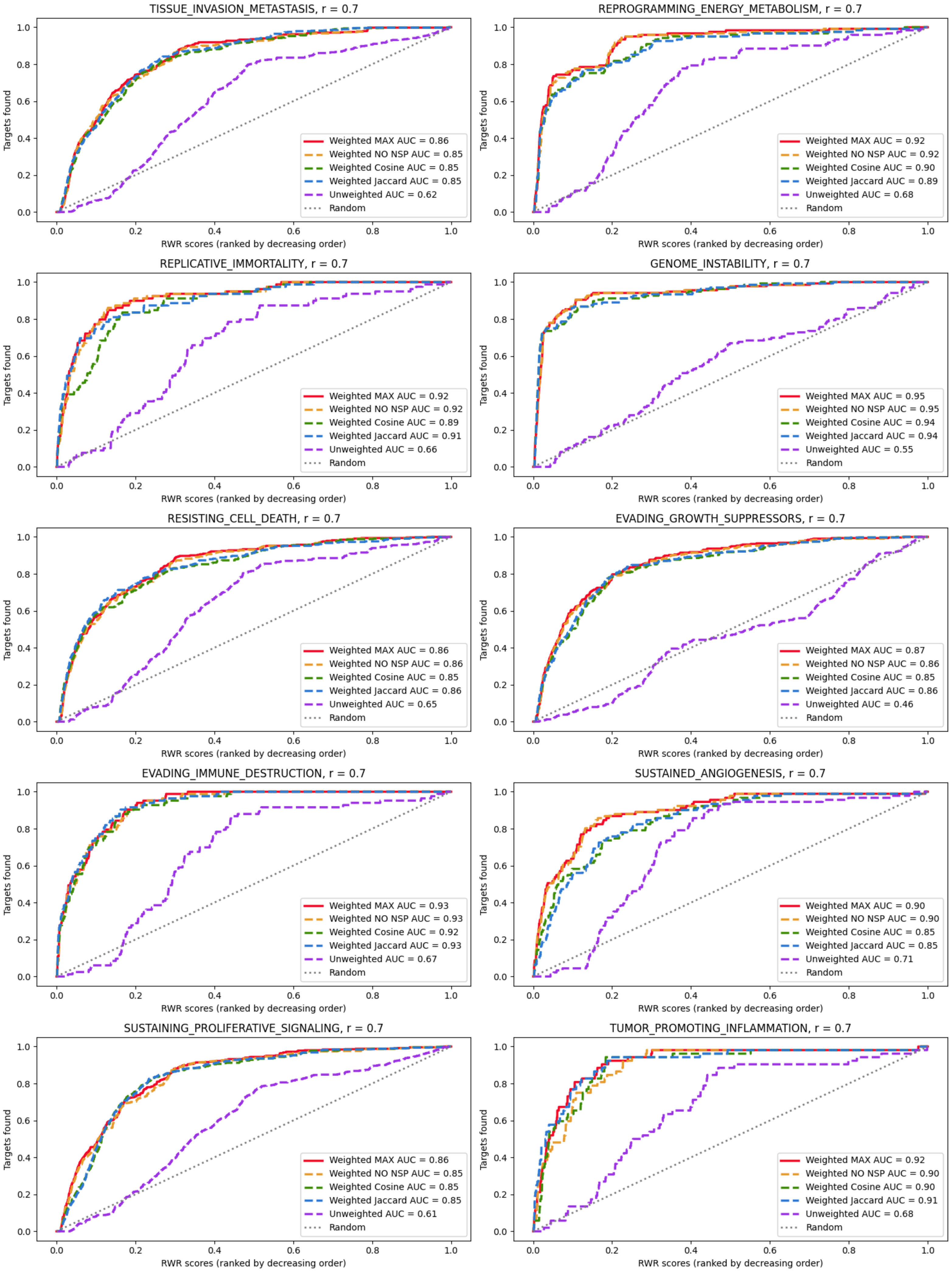
